# PLGA–Soya Lecithin Based Hybrid Nanocomposite for Targeted Topical Delivery of Resveratrol in Psoriasis Management

**DOI:** 10.1101/2025.04.13.648585

**Authors:** Roshan Keshari, Rupali Bagale, Sulagna Rath, Swapnil Raut, Lokesh Kumar Bhatt, Abhijit De, Shamik Sen, Rohit Srivastava

## Abstract

Psoriasis is a recurring autoimmune-driven inflammatory skin disorder characterised by red, scaly, dry skin patches, keratinocyte hyperproliferation and infiltration of inflammatory cells, impacting millions globally. Patients with psoriasis often require repeated treatments with corticosteroids, which are often associated with significant side effects and comorbidities. Therefore, there is a growing need for advanced nanomedicine-based therapeutic approaches. Resveratrol (RSV), a plant-derived polyphenol, has shown promising anti-inflammatory and antioxidant properties for the treatment and management of psoriasis. However, its therapeutic potential is limited due to its poor solubility, low stability, and reduced skin penetration. To overcome these challenges, we have developed dual-carrier conjugated nanoparticles (RSVNPL) composed of soya lecithin and poly(lactic-co-glycolic acid) (PLGA). To further enhance its ease of application and prolong skin residence time, RSVNPL was incorporated into a carbomer 974P-based hydrogel system, forming RSVNPGel. Our formulation demonstrated improved physicochemical properties, including enhanced spreadability, superior skin adhesiveness and stability. *Ex vivo* skin permeation studies confirmed a significantly higher accumulation of RSV in the epidermis and dermis compared to free drugs. *In vivo,* efficacy was investigated using imiquimod (IMQ)-induced psoriatic mouse model, where RSVNPGel treatment led to a notable reduction in epidermal thickness, inflammatory cell infiltration, and key histopathological features associated with psoriasis. Additionally, serum biochemical analysis and organ histology established the biocompatibility and safety of the formulation. Overall, RSVNPGel presents a promising nanocarrier-based strategy for the effective topical management of psoriasis, enhancing RSV delivery and therapeutic efficacy while minimizing systemic exposure.

**Research Highlights:**

- Hybrid nanoparticles composite of soya lecithin and PLGA were developed for RSV delivery.
- The formulation enhanced skin penetration, retention and created a better skin adhesive profile in psoriatic lesions.
- Significant anti-psoriatic effects were achieved in a preclinical model.
- The system showed excellent safety and biocompatibility.

**Graphical Abstract:** 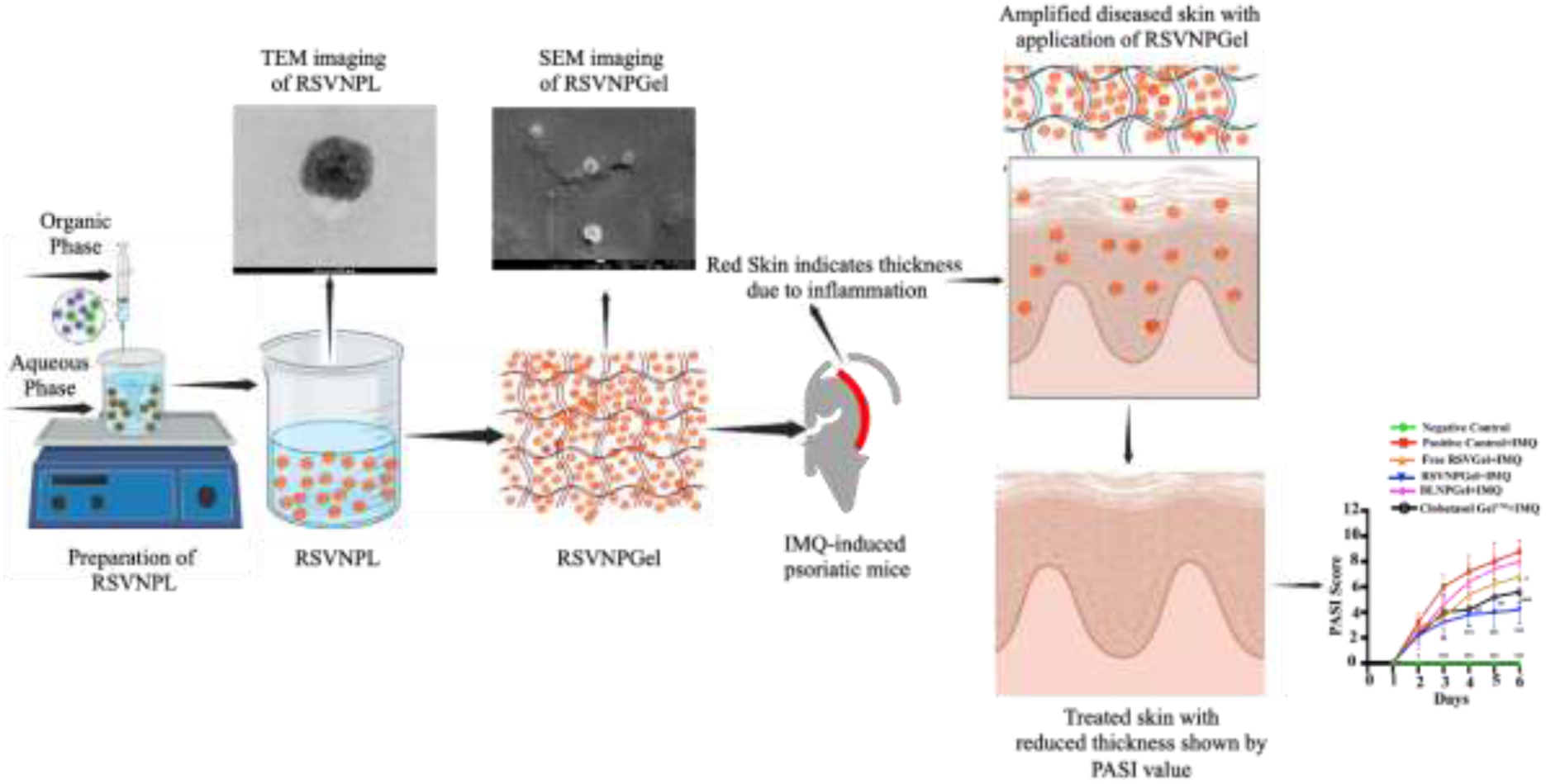

## 1) Introduction

Psoriasis is a chronic autoimmune abnormal skin condition presented with thick, scaly, dry skin, along with abnormal epidermal differentiation, inflammation, infiltration of inflammatory cells and angiogenesis, affecting approximately 2–3% of the global population [1]. It can develop at any age, mainly with two distinct peaks of onset: one typically occurring between the ages of 15 and 20 and another between 55 and 60 years [2]. Although the precise causes and mechanism are not fully understood, it is hypothesised that the IL-23/IL-17 pathway is crucial in driving keratinocyte hyperproliferation and abnormal differentiation, leading to psoriatic lesions. Significant immune cell infiltration has been observed, particularly in the dermis, with proliferating macrophages and T cells contributing to the inflammatory response. Among these, T cells release cytokines that initiate the inflammatory responses, for example, TNF-α, IL-17, and IL-22, further exacerbating disease progression [3][4], [5], [6], [7].

Treatment options for psoriasis may vary based on severity and response to therapy. However, topical treatments are widely used, including corticosteroids, calcipotriol, and combination therapy, for mild to moderate cases [8], [9]. [10]. For more severe cases or when topical treatments are ineffective, systemic drugs such as acitretin, methotrexate, cyclosporine and biologics are prescribed[11]. However, many of these treatments come with significant side effects such as skin irritation, redness, burning and blistering, while long-term use raises concerns such as organ toxicity, immunosuppression, and increased susceptibility to infections. These challenges necessitate exploring alternative, targeted, and safer treatment modalities.

Nanotechnology has emerged as a promising medical platform, significantly improving the delivery of various therapeutic agents, including hydrophobic and hydrophilic drugs, small-molecule drugs, genes, RNAs, and peptides. It also offers promising advancements in improving the therapeutic effectiveness and pharmacokinetics of many drugs[12][13][14]. Lipid-polymer composite nanoparticles have recently gained substantial attention as advanced drug delivery systems that combine the advantages of both lipid and polymeric nanoparticles. These hybrid structures form a polymer core enveloped by a lipid shell, which enhances drug stability, prevents premature leakage, and prolongs systemic circulation time. It offers high drug-loading capacity, controlled release, and precise targeting at tissue, cellular, and molecular levels, making it a highly effective and reliable drug-delivery platform [15], [16],[17],[18]. [19], [20].

Several studies have demonstrated the efficacy of lipid-polymer composite nanoparticles in psoriasis treatment by enhancing targeted drug delivery, improving drug stability, and controlling drug release [21], [22]. For instance, lipid-polymer composite nanoparticles have been utilised to encapsulate the phosphodiesterase-4 inhibitor AN2728, enhancing its targeted delivery to psoriatic skin cells and improving therapeutic outcomes [23]. Similarly, lipid-polymer composite nanoparticles have also been employed to deliver coenzyme Q10 (CoQ10), a potent antioxidant and anti-inflammatory agent for psoriasis treatment. By integrating the benefits of both lipid and polymer components, these nanoparticles provided enhanced drug stability and controlled release [24]. Another promising approach involves encapsulating clobetasol propionate, a corticosteroid inside lipid-polymer composite nanoparticles; these nanoparticles facilitate deeper skin penetration, limited systemic exposure, and better therapeutic outcomes while significantly minimizing the potential for systemic adverse effects [25]. This carrier system can be used as a versatile platform for transporting various hydrophobic drugs like rapamycin for various skin disorders and has the potential for clinical and commercial use [22].

Resveratrol (RSV), a naturally occurring polyphenol, has demonstrated significant therapeutic potential in alleviating psoriasis-related inflammation. Studies indicate that RSV activates the sirtuin 1 (SIRT1) pathway to inhibit aquaporin 3 (AQP-3), a key regulator of keratinocyte proliferation. This mechanism helps mitigate the excessive cell growth characteristic of psoriasis. Additionally, RSV exhibits potent antioxidant and anti-inflammatory properties, further supporting its role in psoriasis management. The topical application of RSV has shown promising effects due to its ability to penetrate the skin effectively, making it an attractive candidate for transdermal drug delivery [26]. Despite its potential, clinical translation is hindered by its poor aqueous solubility, rapid metabolism, and limited bioavailability. To address these limitations, we have developed a lipid-polymer composite nanoparticle system encapsulating RSV. Later, these nanoparticles were incorporated into a hydrogel system for easy application and long-lasting effect. By performing physicochemical characterisation, *in vitro* evaluations, *ex vivo* skin penetration studies, and *in vivo* efficacy assessments, we illustrate the potential of RSV-loaded lipid-polymer composite nanoparticles as a novel, effective, and patient-friendly therapeutic approach for managing psoriasis topically. By utilizing nanotechnology, this formulation offers enhanced drug stability, controlled release, and improved bioavailability, overcoming the challenges associated with conventional psoriasis treatments.

## 2) Materials and methods

Soya lecithin (phosphatidylcholine derived from soybeans), phosphate-buffered saline (PBS, pH 7.4), and fetal bovine serum (FBS) were sourced from HiMedia. The poly(lactic-co-glycolic acid) (PLGA) copolymer, comprising a 50:50 molar ratio of DL-lactide to glycolide with a molecular weight ranging between 7,000 and 17,000 Da, was obtained from Nomisma Healthcare, India. Resveratrol was supplied by Zeta Scientific, India. Tween-80, triethanolamine, and carbopol 974P were procured from Sigma Aldrich, USA. Rhodamine 6G (purity ≥97%) was purchased from Merck KGaA, Germany. A 5% w/w imiquimod cream was procured from Glenmark Pharmaceuticals, India. Throughout all experiments, high-purity deionized water (resistivity of 18.2 MΩ·cm) purified by Milli-Q Plus water purification system (Millipore, USA) was used.

### 2.1 Preparation of resveratrol-loaded lipid polymer composite nanoparticles (RSVNPL)

Resveratrol-loaded lipid-polymer composite nanoparticles were synthesized using the solvent evaporation method via self-assembly of soya lecithin (10 mg/mL) and PLGA (5 mg/mL). Initially, soya lecithin (10 mg/mL) was solubilised in Milli-Q water under gentle heating (∼70°C) with continuous stirring at 500–700 RPM to ensure complete liquefaction. Once fully solubilized, Tween 80 (1%) was incorporated under stirring for uniform dispersion.

Separately, PLGA (5 mg/mL) was suspended in acetone under continuous magnetic stirring (500–700 RPM) to form a clear organic phase. Resveratrol (5 mg/mL) was then gradually added to this PLGA solution while maintaining constant stirring to ensure homogeneity. After preparing both phases, the organic phase was gradually introduced into the aqueous phase at a regulated flow rate of 100 µL/min using a syringe pump (Uni Genetics Instrument Pvt. Ltd.). Simultaneously, the mixture was homogenized at 4000 RPM using a high-speed homogenizer (T25 Ultra-Turrax, IKA, Staufen, Germany) to facilitate nanoparticle formation. To ensure the complete removal of the organic solvent, the resulting formulation was stirred overnight at room temperature (500–700 RPM) on a magnetic stirrer.

Following solvent evaporation, the nanoparticles were collected via centrifugation (Eppendorf Centrifuge 5810 R) at 10,000 RPM for 15 minutes at 4°C. The supernatant was gently discarded, and the resulting pellet was washed three times with Milli-Q water. Finally, the nanoparticles were redispersed in Milli-Q water for subsequent characterization.

### 2.2 Derivation of RSVNPL into a hydrogel (RSVNPGel)

The resveratrol nanoparticle derived hydrogel system (RSVNPGel) was prepared by incorporating carbopol 974P (0.8% w/v) in RSVNPL. To achieve a homogeneous mixture, carbopol 974P was dispersed in RSVNPL and stirred continuously at 800 RPM using a magnetic stirrer. Triethanolamine (6.5 µL/mL) was incorporated into RSVNPGel to maintain the pH to 6.5. Following this, the formulation was homogenized with a high-speed homogenizer until a uniform consistency was achieved.

To ensure the integrity of RSVNPL was maintained upon its conversion into the RSVNPGel, additional characterization techniques, such as scanning electron microscopy (SEM) and texture analysis, were executed to understand the quality and structural properties of the hydrogel system. Some further studies including thermal analysis using differential scanning calorimetry (DSC), molecular structure evaluation using Fourier-transform infrared spectroscopy (FTIR), and crystallinity assessment through X-ray diffraction (XRD) were also performed.

### 2.3 Analysis of Z-average particle size distribution and surface zeta potential

The Z-average particle size distribution, polydispersity index (PDI), and surface zeta potential of RSVNPL were analyzed using dynamic light scattering (DLS) with a (Malvern Zeta sizer Malvern Instrument Ltd., UK) by diluting formulation 100 times with Milli-Q water.

### 2.4 TEM imaging

The structure of RSVNPL was analyzed via TEM imaging by placing a drop of RSVNPL on a carbon-coated copper grid (300 mesh) and allowing adsorption for one minute. Excess liquid was then gently removed using filter paper. To enhance contrast, a drop of 2% (w/v) phosphotungstic acid (PTA) was applied as a negative stain and allowed to dry completely under ambient conditions. Imaging was executed via TEM (FEI Tecnai G2, F30, USA) operating at 300 kV.

### 2.5 Drug Retention Efficiency (DRE)

The DRE of RSVNPL was assessed through the centrifugation method, followed by drug content quantification via HPLC. RSVNPL were subjected to centrifugation (Eppendorf Centrifuge 5810 R) at 10,000 RPM for 15 min. at 4°C. The supernatant were gently removed, and the resulting pellet was rinsed twice with Milli-Q water to discard any non-retained RSV. For RSV quantification, the RSVNPL was diluted tenfold with methanol, and the total RSV retained inside the carrier system was determined through HPLC, as outlined in the corresponding section 2.2.9. The final DRE was calculated using Equation 1

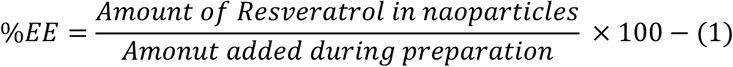

### 2.6 *In vitro* antioxidant assay

2,2-diphenyl-1-picrylhydrazyl (DPPH) free radical scavenging assay was executed to investigate the antioxidant activity of RSVNPL. Briefly, 100 µL of RSVNPL was incorporated within 3.9 mL of DPPH solution (0.1 mM in 95% ethanol). The Absorbance readings were taken at 516 nm at various time intervals such as 5, 10, 20, 40, 60, and 90 minutes. The antioxidant assay of RSVNPL was established using the following:

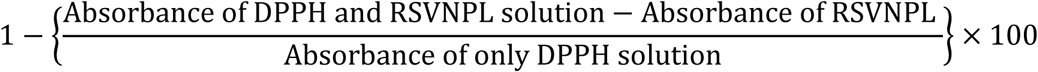

### 2.7 Structural analysis of RSVNPGel via SEM Imaging

SEM imaging was performed to assess the structural integrity of RSVNPL upon incorporation into RSVNPGel. The RSVNPGel sample was carefully placed between two glass coverslips and lyophilized to remove any residual moisture. Once dried, a thin iridium layer was sputter-coated onto the sample to improve electrical conductivity. The surface integrity was then examined with a FEG SEM (FEG-SEM; JEOL JSM-7600F, Japan) operating at a magnification of 50,000× at 10.0 kV and a 4.0 mm working distance for high resolution.

### 2.8 FTIR Analysis

The structural characteristics and functional groups of different formulations were studied using FTIR spectroscopy, including RSVNPGel, Free RSVGel (resveratrol incorporated into carbomer gel), blank nanoparticle gel (BLNPGel), and Free Resveratrol Powder (RSV Powder). For analysis, each sample was carefully placed onto a potassium bromide (KBr) pellet and allowed to dry under an infrared lamp to remove residual moisture. FTIR spectrometer (Bruker, Leipzig, Germany) utilizing a DTGS detector was used to perform spectral measurements using Vertex 80 FTIR spectrometer. Spectra were collected over the 400–4000 cm⁻¹ range, an average of 32 scans per sample with a resolution of 4 cm⁻¹. Post-processing steps, including baseline correction, were carried out using OPUS software.

### 2.9 Rheological behaviour

The rheological characteristics of RSVNPGel were investigated with a rheometer (ARES G2, TA Instruments; MCR 702, Anton Paar, USA). Measurements were performed with a 25 mm parallel plate setup and a 0.5 mm gap. A shear rate of 10 s⁻¹ was applied by rotating the shaft continuously for 3 minutes, followed by a 9-second equilibration period at 25°C. Viscosity was measured at 20 points, averaging three readings. Continuous shear stress measurements recorded 180 data points, varying the shear rate from 0.1 to 100 s⁻¹, with shear stress (mPa) measured at 37°C[27].

### 2.10 Texture Analysis

The textural properties of the RSVNPGel were investigated under compression mode using a CT3-1000 Texture Analyzer (Brookfield Engineering Labs, Inc.) equipped with the TA2/1000 probe. The setup consisted of a cone-shaped male probe for penetrating inside the gel and a corresponding female probe, which contained an RSVNPGel. During the test, the male probe moved toward the female probe at a regulated pretest speed, compressing the gel at 2.0 mm/s until a predetermined interval was attained, with a trigger load set to zero. Upon compression, the probe was withdrawn at the same rate as insertion. Data collection and analysis were performed using TexturePro software[28], [29].

### 2.11 Stability studies of RSVNPGel

Stability studies of RSVNPGel were performed under distinct storage setups, such as RT, 4°C, and accelerated conditions (40°C and 75% RH) for two months. Every month, samples were collected and studied for drug content using visual parameters such as syneresis, consistency, colour change, phase separation, and grittiness. For total drug quantification, 100 mg of RSVNPGel was precisely weighed and dissolved in 900 µL of methanol. The mixture was sonicated in a bath sonicator to facilitate complete drug extraction. Following sonication, the samples were centrifugated at 10,000 RPM for 5 minutes using an (Eppendorf Model-5810 R, Germany). The transparent supernatant solution was collected and analysed for drug quantification using the HPLC method outlined in 2.2.9.

### 2.12 DSC Analysis

Thermal properties exhibited by the different groups, including RSVNP Gel, Free RSV Gel, BLNPGel, and pure RSV powder, were analyzed using a Hitachi Differential Scanning Calorimetry (DSC) 7020 instrument. Approximately 10 mg of the samples from each group were weighed and transferred in an aluminium pan and heated from 30°C-300°C at 10°C/min, with a nitrogen flow of 50 mL/min for accurate thermal analysis. The obtained thermograms were evaluated to determine possible interactions, melting transitions, and the thermal stability of the formulations.

### 2.13 XRD studies

The crystalline or amorphous nature of different groups (RSVNPGel, Free RSVGel, BLNPGel and RSV Powder) was executed via XRD instrument ( PANalytical Empyrean X-ray diffractometer The Netherlands) with a Cu target X-ray tube. The different groups were precisely aligned on the sample holder, and diffraction signal outputs were acquired across a 2θ range of 3° to 40° at 45 kV and 40 mA, ensuring a comprehensive evaluation of the structural attributes of the various groups.

### 2.14 Spreadability Study

The spreadability of RSVNPGel was investigated using the "slip and drag" technique. In brief, 1g of RSVNPGel was positioned between a stationary glass plate and a movable upper glass plate. To achieve uniform distribution, a 500 g weight was placed on the upper plate. Following this, an additional 39g weight was applied to initiate the sliding motion of the upper plate. The total time covered by the upper plate to slide, a distance of 6.03 cm, was recorded. The total spreadability was estimated using the below formula:

S= M* L/T

Where,

S= Spreadability of gel
M = Weight (g) upper plate
L = Plate Distance (cm) travelled
T = Time (s) for the plate to slide 6.03 cm

### 2.15 *In vivo* biocompatibility of RSVNPGel

To evaluate the *in vivo* biocompatibility of the nanoparticles, 250 mg (shaved area of approximately 3.0 cm × 2.5 cm) equivalent of PBS, RSVNPGel, BLNPGel (blank nanoparticle gel), or Free RSVGel was topically applied to the dorsal skin of mice once daily. 72 hours post-treatment, the mice were euthanized, and tissue samples were harvested and preserved (10% neutral buffered formalin) for 24 hours. The samples were processed using paraffin embedding and subsequently sectioned, stained with H&E, and analysed for histopathological study using a Zeiss Axio Imager Z1 microscope (Carl Zeiss, Germany).

### 2.16 *Ex vivo* transdermal penetration and retention study

The penetration profiles of RSVNPGel and free resveratrol gel (Free RSVgel) were investigated via the excised psoriatic skin of Swiss albino mice via Franz diffusion apparatus, based on the methodology presented in Pukale et al. [25]. Briefly, from the excised psoriatic skin, underlying adipose tissue was carefully removed, and the remaining skin was clamped between the administering and receiving compartments of a Franz diffusion apparatus. PBS was prior added to the receiving compartments, and RSVNPGel and Free RSVGel were transferred to the administering compartment for *ex vivo* penetration. The system was continuously stirred at 500–700 RPM and kept at 37°C for 24 hours.

Following incubation, skin samples were removed, rinsed thoroughly three times with PBS to eliminate the adhered drug to the surface, and subsequently dried. The tape-stripping method was employed to selectively remove the stratum corneum layer, discarding the first two strips to remove unabsorbed drugs employing the procedure outlined by Elshall et al.[30]. The next 10 strips were collected and transferred in a container consisting of methanol at 4°C under very slow stirring for 24 hours for complete extraction of RSV.

The remaining tissue was finely minced using a homogenizer, and the resulting homogenate was subjected to gentle stirring at 4°C for 24 hours to facilitate complete extraction. Later on, after incubation, tissue solution was centrifuged, supernatant was collected, and total drug content was analysed via HPLC in accordance with the methodology presented in section 2.2.5.

### 2.2 *In vivo* study using an imiquimod (IMQ)-triggered psoriatic mice model

#### 2.2.1 Psoriatic mice model established using IMQ application

Male BALB/c mice (7–8 weeks old) were procured from the animal facility at the Advanced Centre for Treatment, Research & Education in Cancer (ACTREC), Navi Mumbai, India. The animals were kept at a 14-hour light and dark cycle, a controlled temperature of 22 ± 2°C, and a relative humidity of 50–70% with free access to water and a standard meal. All *in vivo* experiments were performed after receiving the following approval from the Institutional Animal Ethics Committee (IAEC)(Approval No. CPCSEA/IAEC P-33/2023) by following all the instructions strictly decided by the Committee for the Purpose of Control and Supervision of Experiments on Animals (CPCSEA) under the Ministry of Environment and Forests, Government of India.

#### 2.2.2 Investigation of penetration and deposition profile of R6G-based formulations inside the psoriatic skin

*In vivo*, skin penetration and deposition profile of Rhodamine 6G (R6G) labelled formulations were studied in IMQ-induced psoriatic mice ears. R6G@RSVNPGel was prepared following the same protocols used for RSVNPGel. Prior to the experiment, the ears of IMQ-treated mice were gently cleaned, dried, and respective formulations were administrated topically onto the skin and incubated for 24 hours to allow for penetration. After the incubation period, the mice were sacrificed, and ear tissues were excised. Residual formulation on the skin surface was carefully removed with multiple PBS washes. The samples were then imaged with a laser scanning confocal microscope (Carl Zeiss, LSM 780) at 10× magnification, capturing images vertically in a series at 5 µm intervals to understand the penetration depth.

#### 2.2.3 Study of retention and penetration characteristics of nanoparticles via IVIS system

To evaluate the retention and penetration behaviour of RSVNPGel in both healthy and psoriatic mice skin, Rhodamine 6G (R6G)-labelled RSVNPGel (R6G@RSVNPGel) was applied topically to the dorsal skin at a dose of 250 mg (shaved over an area measuring 3.0 cm × 2.5 cm). Fluorescence imaging (ex/em-525/548 nm) was conducted immediately using the *In vivo* imaging system(IVIS) (PerkinElmer, USA) after application at the initial time point to monitor initial distribution. After one hour of incubation, the gel was removed, and the skin was cleaned and imaged again. Subsequently, the formulations were reapplied and incubated for 24 hours. The skin was cleaned and imaged again to check drug retention, which was followed for another 48 and 72 hours.

After IVIS imaging, the mice were euthanized, and key organs such as the heart, kidneys, liver, spleen, and lungs were harvested. These organs were further imaged using the IVIS to assess potential dye retention, providing insights into systemic penetration from the skin.

#### 2.2.4 Investigation of skin adhesion property of RSVNPGel

To evaluate the water resistance and skin adhesive properties of RSVNPGel, approximately 250 mg of R6G@RSVNPGel was topically applied to the shaved dorsal skin of both healthy and psoriatic mice. The surface was gently wiped with a cotton swab after two minutes of application. Any remaining dye was then rinsed with continuous water irrigation. This was followed by fluorescence imaging via IVIS (PerkinElmer, USA) to evaluate formulation retention after washing.

#### 2.2.5 Assessment of treatment effectiveness in an IMQ-triggered psoriasis model

Following a one-week acclimatization period, the mice were randomly assigned into six groups. The dorsal skin of each mouse was carefully shaved over an area measuring 3.0 cm × 2.5 cm in preparation for the study. All animals, with the exception of the negative control group, were given 62.5 mg of commercially available 5% IMQ cream (Imiquad Cream®, Glenmark Pharmaceuticals Ltd., Goa, India) topically on their depilated backs every day for five days in a row. The mice were then assigned to the following treatment groups according to the protocol: the negative control group, which received no topical application of IMQ; the positive control group, which received topical IMQ for five days; the IMQ + BLNPGel group, which received topical IMQ followed by a blank nanoparticle gel (without drug) for five days; the IMQ + RSVNPGel group, which received topical IMQ followed by a resveratrol-based nanoparticle gel for five days; the IMQ + Free RSVGel group, which received topical IMQ followed by a topical free resveratrol formulation in a gelling system for five days; and the Clobetasol^TM^ group, which received topical IMQ followed by a marketed clobetasol formulation for five days. All mice were assessed for scaling, erythema, skin thickness score, and total Psoriasis Area and Severity Index (PASI) scores on days 1, 2, 3, 4, 5, and 6. Skin samples were taken from each mouse using a sterile skin punch, the animals were sacrificed, and the spleen weight was noted at the end of the six-day treatment period. After being gathered, the samples were preserved in 4% formaldehyde for histological examination.

#### 2.2.6 Relative spleen weight to body ratio

The spleen is considered one of the most vital organs connected to the immune system, which is essential for phagocytosis, immunological response, and eliminating infections, foreign antigens, and ageing red blood cells. [31]. Immune cell activation and accelerated proliferation are indicated by an increased spleen weight in relation to body weight. [32]. Post-experiment, mice were sacrificed, their spleens were carefully extracted, and weight was promptly measured to prevent any inaccuracies due to dehydration.

#### 2.2.7 Histopathological analysis

Following euthanasia, dorsal skin tissues were excised and immediately preserved in 4% paraformaldehyde for histopathological study. The tissues were subjected to a dehydration process using a graded ethanol series, after which they were embedded in paraffin blocks. Thin tissue sections were then prepared on glass slides, stained with hematoxylin and eosin (H&E) and imaged using a bright-field microscope (Olympus BX63 upright microscope, cellSens Software, Japan). Several key histological parameters were assessed, including angiogenesis, suprapapillary thinning, inflammatory infiltrates, Kogoj’s pustule formation, hyperkeratosis, parakeratosis, and epidermal hyperplasia. Additionally, a comprehensive total skin damage score was determined to assess the severity of psoriatic lesions.

#### 2.2.8 Assessment of *In Vivo* Drug Safety

To establish the safety profile of the various treatment groups, an *in vivo* cytotoxicity study was performed by collecting primary tissues, including skin, heart, lungs, kidneys, and spleen, from different treatment groups, along with the monitoring of changes in body weight throughout the study period. Furthermore, serum biochemical markers, such as uric acid, alanine aminotransferase (ALT), aspartate aminotransferase (AST), and creatinine, were analysed to understand potential systemic toxicity across the groups.

#### 2.2.9 Apparatus and chromatographic condition

Chromatographic analysis was carried out using a high-performance liquid chromatography (HPLC) system equipped with a Jasco detector 4180 isocratic pump, Jasco 4050 autosampler, and Jasco MD 4015 UV detector was used to quantify the concentration of RSV in RSVNPL and RSVNPGel. An Agilent ZORBAX SB-C18 column (4.6 × 250 mm, 5 µm) was used for separation. The mobile phase comprised acetonitrile and water (50:50, v/v), flowing at 1.0 mL/min. A 20 µL injection volume was used and detected at 307 nm. The method exhibited linearity of (R² = 0.999) within the concentration window of (1–50 µg/mL). Data analysis was carried out using Chrom Nav software.

#### 2.2.10 Statistical Analysis

All results were presented as mean ± standard deviation (SD) or standard error of the mean (SEM). Statistical significance among different groups was determined using repeated measures ANOVA or Student’s t-test, performed via GraphPad Prism (version 8, San Diego, CA, USA). A p-value of *<*0.05 was considered statistically significant. Asterisks indicate p-values in figures (**p *<*0.01,***p *<*0.001,****p *<*0.0001).

## 3. Results and Discussion

### 3.1 Preparation & characterization of RSV nanoparticles (RSVNPL) and RSVNPGel

The formulations were optimized with an emphasis on important factors, including hydrodynamic particle size, surface charge, polydispersity index (PDI) and drug retention efficiency. The dynamic light scattering (DLS) analysis showed that RSVNPL had an average particle size of ∼120 nm **(Fig. 1A)**. A smaller particle size facilitates better skin contact, thereby enhancing drug deposition and penetration. The PDI of 0.214 indicates that nanoparticles are uniformly distributed, while the zeta potential of −31.4 ± 1.6 mV **(Fig. 1B)** suggests good colloidal stability, preventing particle aggregation and ensuring prolonged stability.

**Fig. 1:**
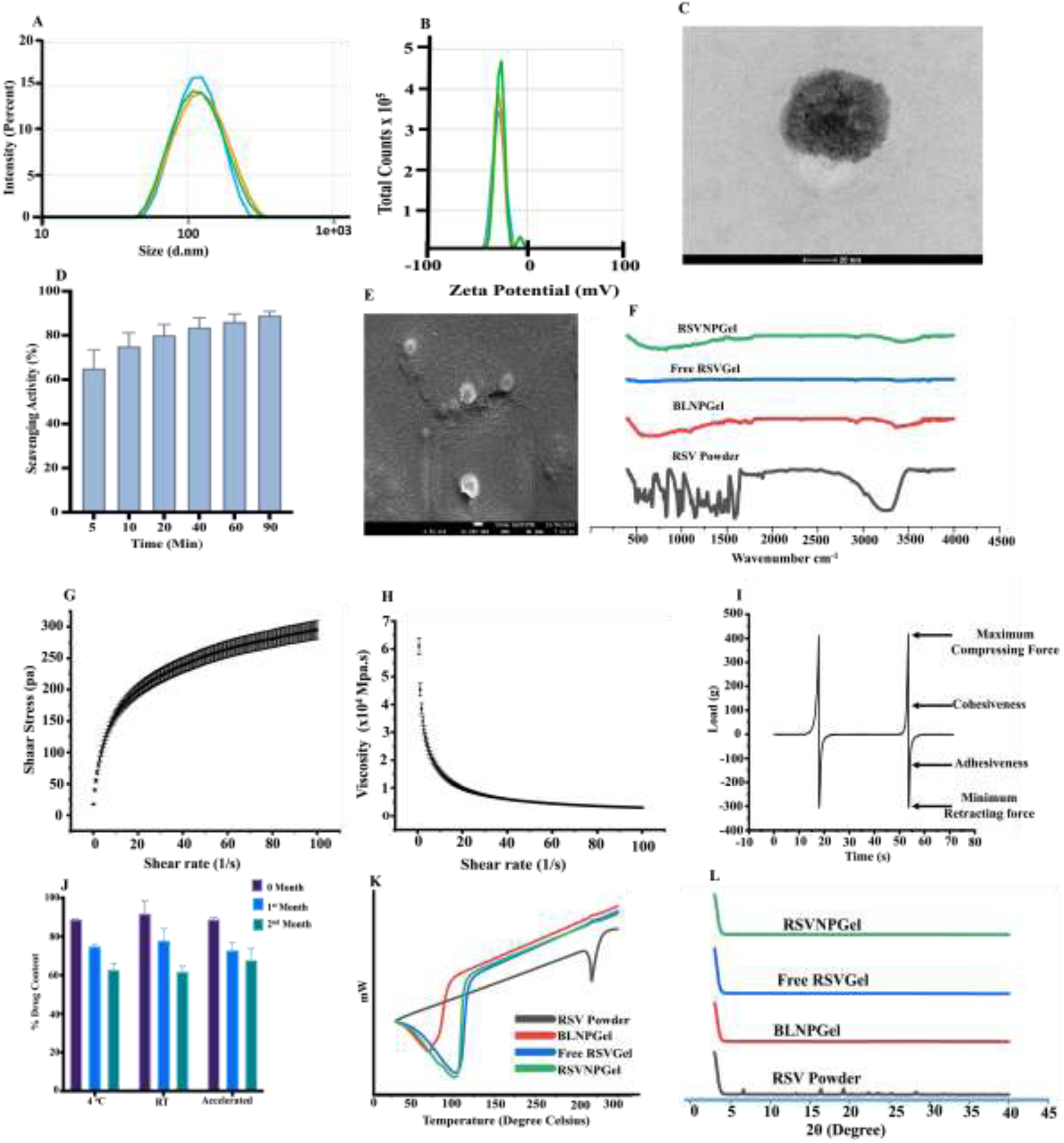
(A) Hydrodynamic particle size distribution of RSVNPL. (B) Zeta potential analysis indicating colloidal stability. (C) The TEM image confirms the spherical morphology and size consistency of RSVNPL with the DLS value. (D) DPPH-based free radical scavenging assay evaluating the antioxidant potential of RSVNPL.(E) SEM image showing RSVNPL successfully embedded within the RSVNPGel matrix. (F) FTIR spectra of various formulations, confirming encapsulation and molecular interactions. (G & H) Rheological analysis: (G) Shear stress vs. shear rate graph demonstrating viscoelastic behaviour and (H) Viscosity vs. shear rate graph illustrating pseudoplastic, shear-thinning properties. (I) Texture profile analysis of RSVNPGel, assessing hardness, adhesiveness, and cohesiveness. (J) The stability data at various storage conditions. (K) DSC thermogram demonstrating the amorphous transition of RSVNPGel compared to free RSV powder. (L) The XRD pattern shows a loss of crystallinity upon encapsulation. All information is shown as mean ± SD (n=3).

TEM imaging **(Fig. 1C)** showed nanoparticles were spherical in shape and ∼100 nm in size, aligning well with the DLS data (Fig. 1A). Furthermore, a drug retention efficiency of ∼54% suggests that more than half of the drug was successfully incorporated into the carrier system. Since RSV is a polyphenolic compound with strong antioxidant properties, a DPPH-based free radical scavenging assay was performed to understand its activity at different time intervals **(Fig. 1D)**. The results showed that RSVNPL exhibited approximately 65% antioxidant activity within 5 minutes of incubation, which progressively increased to 90% by 90 minutes, confirming its potent free-radical scavenging capability. This property is particularly beneficial for psoriasis patients, where oxidative stress plays a major role in disease progression.

Upon incorporation of RSVNPL into RSVNPGel, SEM imaging **(Fig. 1E)** was performed to confirm that the nanoparticles retained their integrity within the hydrogel system when incorporated. The images confirmed that RSVNPL particles were successfully embedded within the hydrogel system without compromising their structural integrity, which is essential for enhancing drug penetration, preventing premature drug leakage, and ensuring sustained release upon skin application.

To confirm the efficient encapsulation of RSV within the nanoparticle system and its successful integration into the hydrogel matrix, FTIR imaging was performed on RSVNPGel, Free RSVGel (i.e., RSV in carbomer gel), blank nanoparticle Gel (BLNPGel), and free resveratrol powder (RSV Powder) **(Fig. 1F)**. The characteristic O-H stretching vibrations were observed between 3300–3500 cm⁻¹, with peaks at 1680–1710 cm⁻¹ and 1600– 1620 cm⁻¹ corresponding to carbonyl (C=O) and alkene functional groups, respectively, confirming the presence of RSV. Additionally, peaks at 1750–1780 cm⁻¹ and 2800–3000 cm⁻¹ confirmed the presence of PLGA, while soya lecithin components gave rise to peaks at 1500–1550 cm⁻¹ and 1630–1680 cm⁻¹. Notably, RSVNPGel did not exhibit the distinct peaks of free RSV, suggesting successful encapsulation and integration into the hydrogel matrix.

The rheological properties of RSVNPGel are significant in determining its spreadability and skin residence time at the application site. While the viscosity vs. shear rate graph **(Fig. 1H)** showed a decrease in viscosity with increasing shear rate, the shear stress vs. shear rate graph **(Fig. 1G)** validated the viscoelastic behaviour of the gel. These findings indicate that RSVNPGel exhibits pseudoplastic (shear-thinning) behaviour, making it suitable for topical applications, as it allows easy application while maintaining its consistency on the skin.

A texture analysis was performed to evaluate cohesiveness, adhesiveness and hardness (**Fig. 1I)**. The hardness of the gel, representing the maximum force required for deformation, was ∼410 g, confirming its polymeric rigidity and mechanical stability. The adhesiveness, which reflects the gel’s ability to stick to the skin, was measured to be ∼5.5 mJ; the adhesive force was estimated to be 306 mJ, indicating strong skin adherence, which is ideal for prolonged therapeutic effects.

Additionally, the slip and drag technique was used to assess the spreadability of RSVNPGel which showed a value of 5.88 ± 0.14 g cm/sec, confirming smooth application with minimal force, ensuring uniform drug distribution across the skin.

The stability of RSVNPGel was assessed over a two-month period under different storage conditions, including room temperature (RT), 4°C, and accelerated conditions (45°C/75% RH) **(Fig. 1J)**. The formulation showed acceptable stability under 4°C and ambient conditions for up to one month, with noticeable degradation and variability observed across all storage conditions by the second month, indicating time-dependent degradation. Additionally, no changes in grittiness, syneresis, odour or visual appearance, suggesting no degradation during storage, further confirming the stability and integrity of the formulation.

To investigate the physical state of RSV within the formulation, DSC analysis was performed **(Fig. 1K)**. The DSC thermogram of free RSV powder displayed a distinct endothermic peak near 250°C, providing evidence of crystallinity. However, these sharp peaks were unavailable in RSVNPGel, suggesting successful entrapment and transformation to an amorphous form. This transformation is beneficial as it can enhance solubility, bioavailability and overall drug stability. To know more about the crystalline characteristics of RSVNPGel, XRD study was performed **(Fig. 1L)**. The XRD analysis of RSV powder revealed a crystalline nature, while RSVNPGel showed amorphous characteristics, suggesting successful encapsulation and enhanced drug stability due to the gel matrix. Overall, the preparation and characterisation of RSVNPL and RSVNPGel demonstrated successful nanoparticle formulation, efficient drug encapsulation, excellent skin adhesion, rheological suitability, and prolonged stability.

### 3.2 Enhanced skin delivery of RSVNPGel for targeted psoriasis treatment

Franz diffusion apparatus was used to understand *ex vivo* transdermal skin penetration and deposition of RSVNPGel and Free RSVGel in excised psoriatic mice skin. Compared to Free RSVGel, RSVNPGel indicated a substantially increased accumulation of RSV in the epidermis and dermis, with penetration levels roughly increased by ∼2.5 times and ∼1.5 times, respectively. (**Fig. 2A & B**). This enhanced permeation can be attributed to RSVNPGel exerting an occlusive action by forming a thin layer on the surface of the skin, which helps minimize transepidermal water loss and improves, thereby, drug retention.

**Fig. 2:**
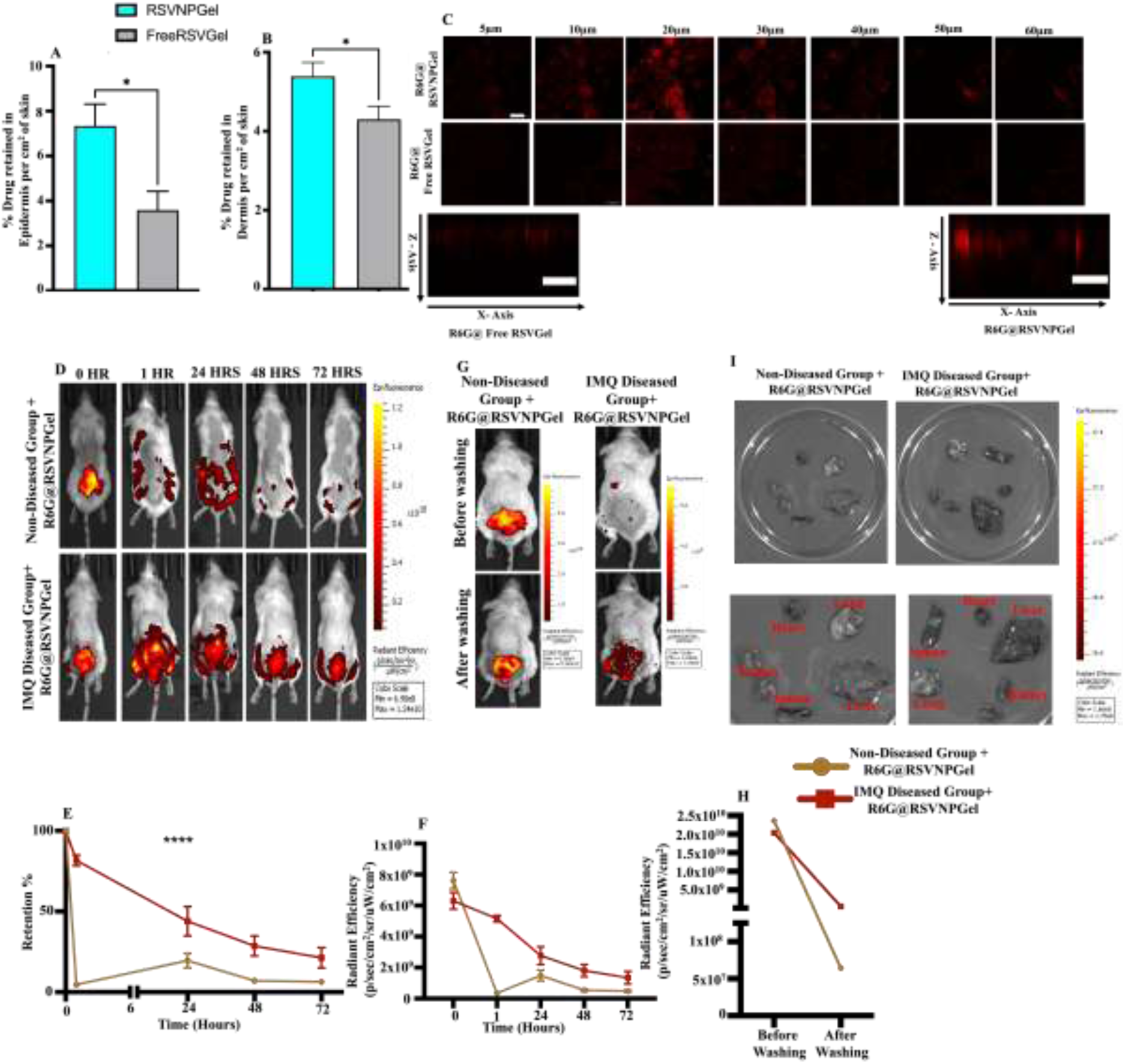
(A, B) *Ex vivo* skin permeation and deposition of RSVNPGel and free RSVGel in epidermis and dermis of excised psoriatic mouse skin. (C) CLSM imaging of psoriatic ear skin after 24 hours of treatment with R6G@RSVNPGel and R6G@FreeRSVGel. (D, E&F) IVIS imaging analysis of skin deposition and excretion profiles of R6G@RSVNPGel over three days in diseased and non-diseased mice and their retention and radiant efficiency profile. (G&H) Skin adhesion study of R6G@RSVNPGel in diseased and non-diseased mice and radiant efficiency (I) IVIS imaging of major organs (lungs, heart, kidneys, spleen, and liver) to assess the systemic accumulation of RSVNPGel. (Scale bar: 132 μm).

To further measure the penetration depth of RSVNPGel, the formulation was labelled with rhodamine 6G, a fluorescent dye, and its permeation was tracked in psoriatic skin using confocal laser scanning microscopy and further compared with R6G@RSVNPGel and R6G@FreeRSVGel. After 24 hours of application, CLSM imaging of psoriatic ear skin showed a higher fluorescence intensity in the R6G@RSVNPGel group, diffusing through the layers up to a depth of 60 μm **(Fig. 2C)**. This diffusion enabled RSVNPL to diffuse through the layers of psoriatic skin, reaching to the epidermis, where it was readily taken up by keratinocytes and infiltrating immune cells, thereby enhancing its therapeutic efficacy. Additionally, the X-Z axis orthogonal skin visualisation demonstrated a uniform distribution of RSVNPGel, providing further evidence of its improved penetration ability.

The skin penetration and deposition profile of RSVNPGel were further assessed using an IVIS imaging system. Comparative skin retention and elimination profiles of R6G@RSVNPGel were analysed in both diseased and non-diseased mice. After 24 hours post, the psoriatic group treated with R6G@RSVNPGel exhibited a higher fluorescence intensity than the healthy control group treated with the same formulation. By day three, a reduction in fluorescence signal was observed, suggesting gradual clearance of nanoparticles from the skin **(Fig. 2D)** and the percentage of dye retention and radiant efficiency was also measured **(Fig. 2E&F)**. This decrease was more pronounced in healthy skin, likely due to the lack of strong adhesion of nanoparticles. These findings suggest that R6G@RSVNPGel selectively penetrates psoriatic skin and remains retained for longer, potentially enhancing therapeutic efficacy while minimizing exposure to healthy skin.

To evaluate the skin adhesive property of R6G@RSVNPGel, the formulation was topically applied on the dorsal skin of both psoriatic and healthy mice and subjected to 2 min incubation. This was followed by rinsing with water and gentle wiping with cotton. IVIS imaging revealed a stronger fluorescence signal in psoriatic skin than in healthy skin, indicating enhanced retention in psoriatic skin **(Fig. 2G)** and radiant efficiency **(Fig. 2H)**. This suggests that R6G@RSVNPGel did not adhere well to intact, healthy skin, as it was restricted to the superficial layers of the stratum corneum. However, in psoriatic skin, where the barrier is compromised, and the skin is also rough, R6G@RSVNPGel demonstrated strong adhesion and deeper penetration, highlighting it as a targeted topical treatment for psoriasis.

To evaluate the systemic accumulation of RSVNPGel, major organs, including the lungs, heart, kidneys, spleen, and liver, were isolated from both diseased and non-diseased mice and imaged with an IVIS imaging system **(Fig. 2I).** No fluorescence intensity was detected in any of these organs, confirming RSVNPGel exhibited localized skin retention with minimal systemic absorption, reducing potential off-target effects. Collectively, these findings suggest that R6G@RSVNPGel can easily penetrate deeper skin layers, thereby exerting its therapeutic effects without any detectable accumulation in major organs.

### 3.3 RSVNPGel demonstrates therapeutic efficacy in an IMQ-induced psoriatic mice model

An imiquimod (IMQ)-induced psoriatic mice model was used to evaluate the therapeutic potential of RSVNPGel. This model closely mimics the pathological features of human psoriasis, such as epidermal thickening, scaling and formation of skin patches. As a result, it is widely accepted as a reliable preclinical model for psoriasis treatment. **Fig. 3A** illustrates the experimental timeline, detailing the application of IMQ and various formulations administered daily at 8-hour intervals. **Fig. 3B** illustrates representative images of mice captured from the first day to the day of sacrifice, visually documenting the progression of disease progression and treatment response.

**Fig. 3:**
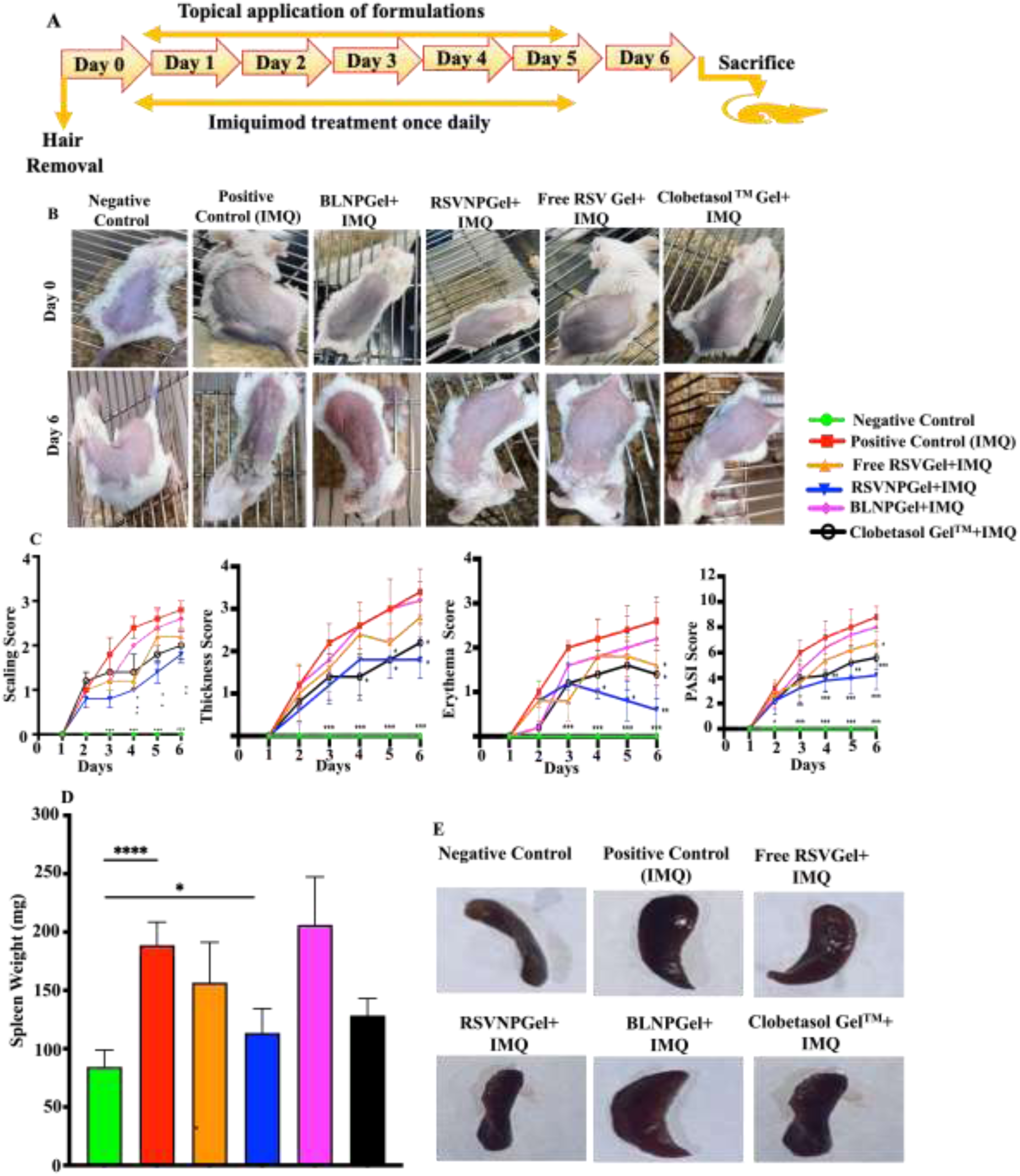
(A) Schematic representation of the experimental timeline, illustrating the induction of psoriasis using imiquimod (IMQ) and the administration of different formulations at 8-hour daily intervals. (B) Representative images showing the progression of psoriasis and treatment response in mice from day 1 to the day of sacrifice. (C) Changes in PASI score over six days demonstrate the therapeutic efficacy of various treatments. (D&E) Spleen weight analysis and their imaging, highlighting IMQ-induced splenomegaly as a marker of systemic inflammation.

The severity of psoriasis and the therapeutic benefit of the formulations were studied via the PASI score. Following the topical application of IMQ for five consecutive days, mice started to develop characteristic psoriatic-like skin lesions, including scaling, erythema and increased back skin thickness **(Fig. 3C)**. In the positive control mice (IMQ-only group), these symptoms progressively increased from day 1 to day 6 with PASI score of ∼9 by day six. However, therapeutic intervention significantly alleviated psoriasis symptoms among the various treatment groups. IMQ **+** RSVNPGel demonstrated the highest therapeutic efficacy, significantly reducing the PASI score to 4.2. In contrast, IMQ + free RSV gel treatment led to a more moderate reduction, lowering the PASI score to 6.8. Meanwhile, IMQ + Blank Gel showed minimal improvement, with a PASI score of 8.0, indicating a limited therapeutic benefit. These findings highlight the superior effectiveness of RSVNPGel in alleviating psoriasis symptoms compared to other treatment groups.

Furthermore, IMQ treatment led to splenomegaly, a hallmark of systemic inflammation, with spleen weight significantly increasing to ∼206 mg in the IMQ group, compared to∼ 84 mg in the negative control group. Notably, the RSVNPGel group exhibited the lowest spleen weight at ∼113.5 mg, closely resembling the negative control **(Fig. D&E)**. These results further support the therapeutic efficacy of RSVNPGel, demonstrating its potential to effectively alleviate psoriasis while minimizing systemic inflammation.

### 3.4 Histological Examination of Psoriatic Skin: Therapeutic Impact of RSVNPGel Treatment

The histological examination of back skin is presented in **Fig. 4**. Parakeratosis, hyperkeratosis, inflammatory cell infiltration, epidermal hyperplasia, capillary proliferation (neo-angiogenesis), suprapapillary thinning, microabscess development, and Kogoj’s pustules were among the important psoriatic skin parameters that were noted. The degree of skin injury in each treatment group was also evaluated using a thorough scoring system. Among them, the RSVNPGel-treated group showed the lowest overall skin damage score **(Table 1)**. Notably, RSVNPGel demonstrated a significant improvement in restoring skin integrity compared to other treatment groups, yielding better therapeutic outcomes than the other formulations.

**Fig. 4.**
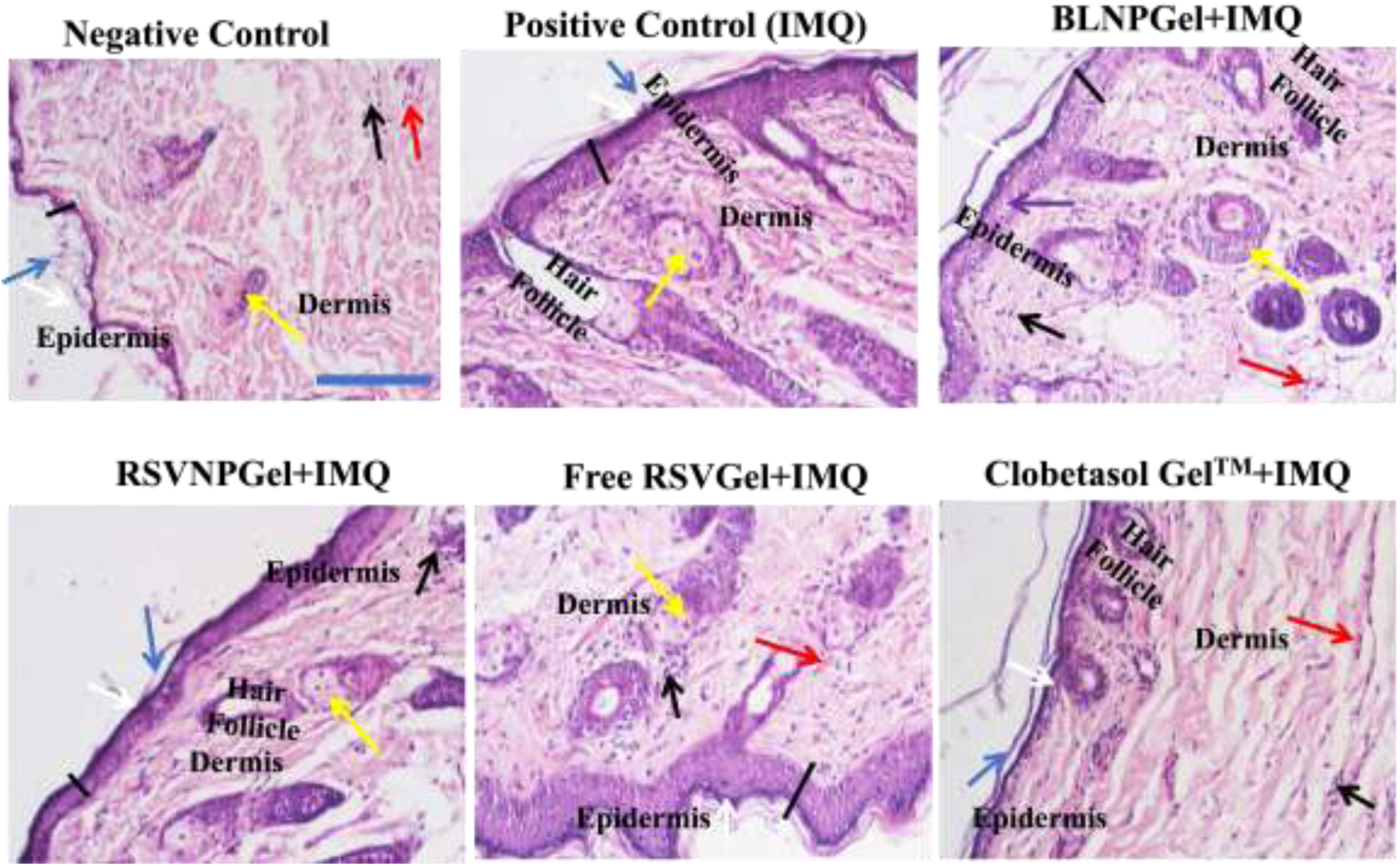
Hematoxylin and eosin (H&E) stained histological assessment of psoriatic back skin following different treatments. Magnification-10X and scale bar:10μm) **Keys: Parakeratosis (Blue Arrow), Hyperkeratosis (White Arrow), Inflammatory Cells (Black Arrow), Neo-Angiogenesis (Red Arrow), Epidermal Thickness (Black Line), Sebaceous Gland (Yellow Arrow), and Epidermal Edema/Microabscess (Violet Arrow)**

**Table-1.**
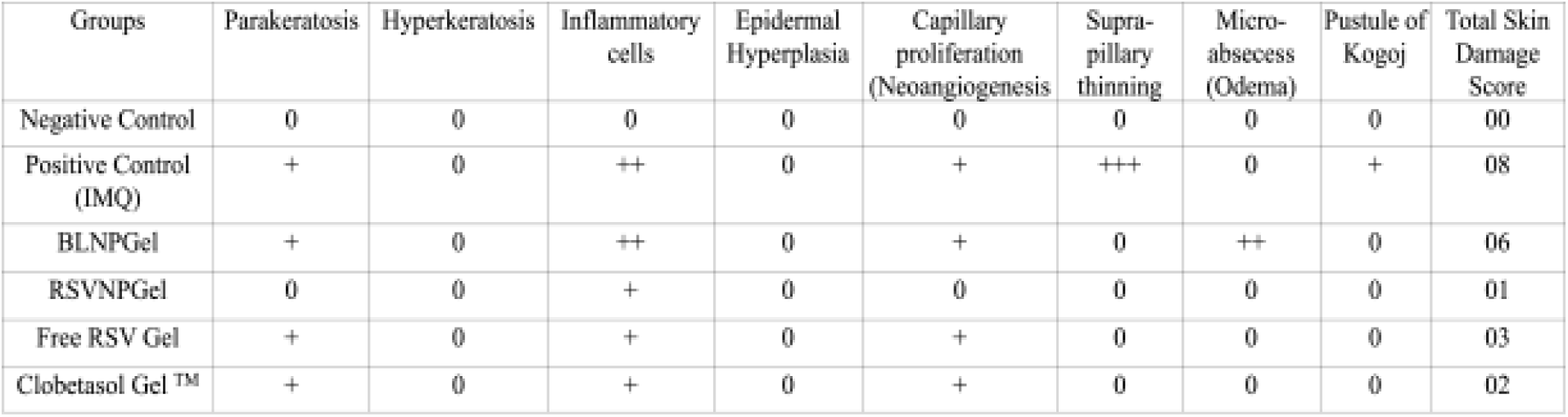
Quantitative scoring of skin damage revealed that the RSVNPGel treated group exhibited the lowest overall skin damage score, indicating superior therapeutic efficacy. (Scoring: + Indicate Mild, ++ Indicate Moderate; +++ indicate Severe)

| Groups | Parakeratosis | Hyperkeratosis | Inflammatory cells | Epidermal Hyperplasia | Capillary proliferation (Neoangiogenesis) | Supra-pillary thinning | Micro-abscess (Odema) | Pustule of Kogoj | Total Skin Damage Score |
| --- | --- | --- | --- | --- | --- | --- | --- | --- | --- |
| Negative Control | 0 | 0 | 0 | 0 | 0 | 0 | 0 | 0 | 00 |
| Positive Control (IMQ) | + | 0 | ++ | 0 | + | +++ | 0 | + | 08 |
| BLNPGel | + | 0 | ++ | 0 | + | 0 | ++ | 0 | 06 |
| RSVNPGel | 0 | 0 | + | 0 | 0 | 0 | 0 | 0 | 01 |
| Free RSV Gel | + | 0 | + | 0 | + | 0 | 0 | 0 | 03 |
| Clobetasol Gel <sup>TM</sup> | + | 0 | + | 0 | + | 0 | 0 | 0 | 02 |

### 3.5 *In vivo* Drug Safety Assessment

To assess the possible toxic effects *in vivo*, a cytotoxicity study was performed in Swiss albino mice over a time period of 72 hours with PBS as the negative control, followed by H&E staining. Interestingly, none of the *in vivo* treatment groups showed any potential toxicity when evaluated against the PBS control **(Fig. 5A)**.

**Figure 5.**
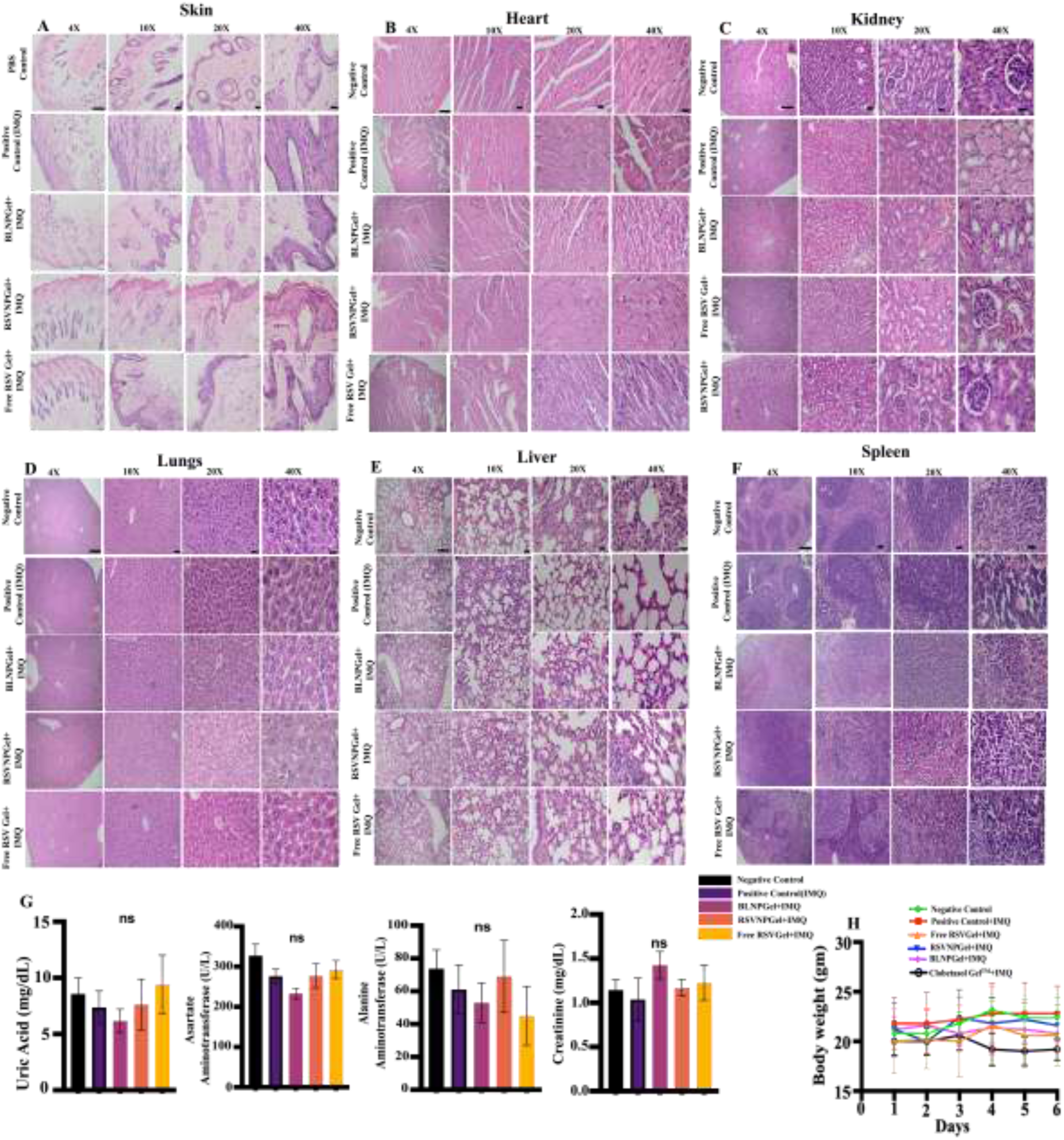
*In vivo* toxicity assessment of the prepared formulation. (A) Representative H&E-*in vivo* cytotoxicity of Swiss albino mice skin treated with the formulation for 72 hours Scale bar. (B–F) Histopathological analysis of the heart, kidney, lungs, liver, and spleen at various magnifications. (G) Serum biochemical analysis of key metabolic markers (H) Body weight monitoring throughout the study. (Scale bar: 4X: 200 μm, 10X: 50μm, 20X&40X: 20μm.

Similarly, post-study, key organs such as the heart **(Fig. 5B)**, kidney (**Fig. 5C),** lungs **(Fig. 5D)**, liver **(Fig. 5E)** and spleen **(Fig. 5F)** were harvested for H&E staining and imaged at various magnifications, i.e. (4X, 10X, 20X, and 40X) to understand the potential adverse effects. No significant histopathological abnormalities, either major or minor, were detected in any of the organs compared to the positive control, further confirming the biocompatibility and safety of the particles for therapeutic use.

Additionally, to assess any metabolic and physiological changes in vital organs, particularly the liver and kidneys, serum biochemical assays **(Fig. 5G)** were carried out. Key biochemical markers such as aspartate aminotransferase (AST), alanine aminotransferase (ALT), uric acid, and creatinine (CRE) were analysed, which suggested no significant variations were observed across the treatment groups, further validating the non-toxic nature of the formulation.

Furthermore, body weight measurements were consistently tracked throughout the experiment duration **(Fig. 5H)**, and no notable differences were detected across groups, suggesting that the treatment did not induce systemic toxicity.

The comprehensive *in vivo* toxicity assessment showed that the prepared formulation showed excellent biocompatibility with no observable systemic or organ-specific toxicity.

Histopathological evaluation and serum biochemical analysis further confirmed its safety, while stable body weight measurements indicated no adverse physiological effects. These findings support the potential of the formulation for safe therapeutic applications.

## 4 Discussion

Psoriasis is a systemic immune-driven inflammatory skin disorder marked by keratinocyte hyperproliferation, aberrant immune cell infiltration, the presence of dense inflammatory cells, as well as an excess of important inflammatory cytokines such as IL-17, IL-10, and IL-23. Managing and treating psoriasis always remains a significant clinical challenge, as most available therapies, such as systemic, topical and biologics, often face many limitations, including poor skin penetration, systemic side effects, dose dumping from frequent high dosing, and reduced efficacy with long-term use. These challenges often lead to poor patient compliance and unsatisfactory treatment outcomes. To address these issues of treatment modalities, there is a growing need for advanced nanotechnology based therapeutic approaches that can effectively ameliorate psoriatic symptoms while reducing their associated adverse effects with available treatments. In this regard, the current work effectively illustrated the therapeutic potential of resveratrol-loaded lipid-polymer composites nanoparticles (RSVNPL), which were then embedded inside a carbopol 974P-based hydrogel (RSVNPGel) for the treating and managing inflammation associated with psoriasis. It was strategically developed as a next-generation nanocarrier designed to enhance the targeted delivery of resveratrol directly to psoriatic lesions while limiting systemic absorption, thereby improving treatment efficacy and safety.

The physicochemical characterization of RSVNPL confirmed the development of an optimized nanoparticulate system with a narrow size distribution of ∼120 nm. Previous studies have indicated that nanoparticles with smaller particle sizes can significantly enhance surface contact with the skin, leading to improved drug penetration and deposition. This effect is particularly beneficial in inflammatory skin conditions such as psoriasis and atopic dermatitis, where the skin barrier is already compromised, allowing nanoparticles to penetrate more effectively, deeply inside the skin layer and deliver therapeutic agents [33]. Zeta potential is considered a key parameter in nanoparticle characterization since it represents the surface charge on the nanoparticles and is crucial in predicting the stability of colloidal systems; a higher zeta potential, either negative or positive (typically above ±30 mV), contributes to enhanced particle stability by generating electrostatic repulsion between particles. This repulsion prevents particle aggregation and supports long-term dispersion stability [34]. The zeta potential of the prepared nanoparticles determined in this study was – 31.4 ± 1.6 mV, indicating a high negative surface charge. This value suggests better colloidal stability, reducing the risk of particle aggregation during storage and ensuring the sustained integrity of the formulation over time. The narrow and consistent size distribution of the nanoparticles was indicated by the PDI, which was 0.21. A PDI value below 0.3 generally reflects a homogeneous population [35], and in the presented study, approximately 80% of the particles were uniformly distributed. Such uniformity is crucial as it ensures consistent and predictable drug distribution to improve reliable therapeutic performance. The total drug retention efficiency inside the carrier system was 54.2%, indicating that approximately half of the drug was successfully encapsulated within the carrier system. This level of encapsulation is considered satisfactory for nanoparticle-based topical formulations, as it ensures an adequate drug payload while maintaining the structural integrity and stability of the delivery system.

The antioxidant study through the DPPH assay showed that RSVNPL possessed significant free radical scavenging activity, a key attribute considering the critical role oxidative stress plays in psoriasis pathogenesis. By reducing oxidative stress, RSVNPGel reduces inflammation and protects keratinocytes from further damage, potentially reducing psoriasis progression to a major extent.

The RSVNPL was further incorporated into a carbopol 974P-based hydrogel matrix to facilitate ease of topical application and to prolong skin residence time upon administration, and SEM imaging was used to confirm its structural integrity, which was intact, and no observable morphological changes were observed. Further studies showed hydrogel exhibited ideal viscosity, spreadability, and adhesiveness, critical properties for ensuring uniform application and extended retention on inflamed psoriatic skin, as supported by previous literature reports[36]. Furthermore, FTIR, XRD, and DSC data provided additional evidence of successful drug encapsulation within the nanoparticles, consistent with findings reported in the literature, thereby ensuring the stability and compatibility of the formulation components[22].

Similarly, the stability of RSVNPGel was studied over two months, which showed a decline in stability across all storage conditions by the second month, suggesting a time-dependent degradation of the formulation. While acceptable stability was maintained for up to one month under refrigerated and room temperature conditions, the loss of integrity beyond this point highlights the need for further optimization, such as the incorporation of stabilizers or advanced preservation strategies, to enhance long-term storage potential.

One of the major obstacles in the topical treatment of psoriasis is overcoming the thickened, hyperkeratotic skin barrier, which significantly reduces the penetration of therapeutic agents into the deeper skin layers. This barrier not only restricts effective drug delivery but also requires higher dosing frequencies, increasing the risk of dose dumping and directly reducing therapeutic efficacy. In the present study, RSVNPGel successfully addressed these limitations, as demonstrated by Franz diffusion studies. Compared to the free resveratrol gel, RSVNPGel achieved approximately 2.5 and 1.5 times higher accumulation of RSV in the epidermis and dermis. This might be attributed to the occlusive nature of the hydrogel, which enhances skin hydration and reduces transepidermal water loss, allowing them to navigate through the compromised psoriatic skin barrier. Moreover, CLSM imaging further confirmed the superior penetrability of rhodamine 6G-labeled RSVNPGel, diffusing up to 60 μm and displaying an evenly spread across the psoriatic skin layers.

The skin penetration, deposition and retention behaviour of RSVNPGel were further validated through IVIS imaging techniques, which showed higher fluorescence intensity and prolonged retention of R6G@RSVNPGel in psoriatic skin compared to healthy skin after 24 hours of incubation, with gradual clearance from the body, was observed by day three. This might be due to the compromised barrier and stronger adhesion of the nanoparticles, as confirmed by skin adhesion studies, which showed superior binding to psoriatic skin over healthy tissue. Interestingly, no fluorescence signals were detected in major organs, indicating no or very minimal non-detecting systemic absorption and further confirming the localised action of RSVNPGel, which minimizes off-target effects while maximizing therapeutic efficacy for psoriasis treatment.

The therapeutic potential of RSVNPGel was understood with the application of IMQ-induced psoriatic mice model, a gold standard method for inducing psoriasis models. This model closely replicates key features of human psoriasis, including epidermal hyperplasia, immune cell infiltration and elevated proinflammatory cytokines. Upon IMQ application, mice developed psoriatic lesions with a PASI score reached approximately 9 by day 6. The topical application of RSVNPGel treatment significantly reduced the PASI score to approximately 4.2, confirming its superior efficacy in alleviating psoriasis symptoms. Additionally, RSVNPGel effectively reduced spleen weight, approximately 113.5mg, comparable to the negative control, indicating its role in minimizing systemic inflammation alongside local skin improvement. These findings align with the previous literature demonstrating the role of resveratrol in psoriasis treatment[37], [38], [39]. These results confirm RSVNPGel as a promising therapeutic strategy for managing and treating psoriasis with enhanced local efficacy and reduced systemic immune activation.

The histopathological study further validated and confirmed the therapeutic efficacy of RSVNPGel. Treated skin showed a remarkable reduction in typical psoriatic features such as parakeratosis, hyperkeratosis, microabscess formation, and inflammatory infiltration, which indicates effective restoration compared to the architecture of normal skin and suppression of local inflammation.

Various experiments were executed to understand the safety profile of the developed formulation, such as a comprehensive *in vivo* toxicity study, which confirmed the better biocompatibility of RSVNPGel. Serum biochemical indicators (AST, ALT, uric acid, and creatinine) remained within normal ranges across all treatment groups, and vital organs such as the heart, kidney, lungs, liver, and spleen showed no histopathological abnormalities.

Additionally, stable body weight throughout the treatment duration and the absence of systemic toxicity further validated the safety of the formulation, supporting its potential for safe and localized therapeutic application in psoriasis.

These results collectively support the idea that the RSVNPGel not only improves the topical bioavailability of resveratrol but also offers superior therapeutic outcomes with multimodal action such as antioxidant activity, anti-inflammatory activity, enhanced skin penetration and adhesion profile, and reduced PASI score. Compared to conventional treatments, including corticosteroids, RSVNPGel presented a safer, non-steroidal alternative that minimizes side effects while maintaining efficacy.

However, while the results of this preclinical study are promising, further research should focus on detailed pharmacokinetic and dermatokinetic studies to thoroughly understand the behaviour of the developed formulations in the skin and systemic circulation. Additionally, long-term *in vivo* studies and clinical trials should be performed, which are essential to fully establish the safety, efficacy, and translational potential of RSVNPGel for clinical use in humans.

## 5 Conclusions

To sum up, based on the presented data, RSVNPL has excellent potential as an advanced hybrid nanomedicine-based strategy for the treatment and management of psoriatic-like skin inflammations. The formulation exhibited excellent physiochemical properties, long-term stability, deeper, improved skin penetration, retention, deposition, and therapeutic efficacy. Histological imaging further confirms the restoration of normal skin architecture. Various experiments, such as histological examination of various organs and unchanged body weight throughout the experiment and biochemical analysis, confirm the safety of the developed formulations. Collectively, these results supported the potential clinical translation of RSVNPGel as an effective, targeted, and localized therapeutic approach for treating and managing psoriasis and related dermatological disorders like skin acne, vitiligo and atopic dermatitis.

## 6 Acknowledgement

The author acknowledges SAIF (Sophisticated Analytical Instrument Facility) and IRCC (Industrial Research & Consultancy Centre) at the Indian Institute of Technology Bombay, India, for providing instrumental support. Roshan Keshari also acknowledges the DAAD (German Student Exchange Programme), Govt. of Germany, for a research fellowship. I would also like to thank Ms. Ankana De, affiliated at the School of Biological Sciences, Indian Association for the Cultivation of Science, Jadavpur, Kolkata-700032, summer Intern at NanoBios lab to help with the IVIS imaging experiment. Graphical abstract (Created with BioRender.com).

## 7 CRediT authorship contribution statement

**RK-** Conceptualization, methodology, execution, investigation, analyzing experiments, literature survey, writing original draft, review & editing. **RB** -Very initial particle characterization, review and editing. **LKB&SR**-*In vivo* studies. **SR&AD-**Histology slide preparation, interpretation and IVIS imaging. **SS&RS**-Conceptualization, Supervision, Investigation, Project administration, Funding acquisition, Resources, review & editing of the manuscript.

## 9. Data availability

Data will be made available on request.

